# Genome features of common vetch (*Vicia sativa*) in natural habitats

**DOI:** 10.1101/2021.03.09.434686

**Authors:** Kenta Shirasawa, Shunichi Kosugi, Kazuhiro Sasaki, Andrea Ghelfi, Koei Okazaki, Atsushi Toyoda, Hideki Hirakawa, Sachiko Isobe

**Author notes:** RIKEN, Yokohama, Kanagawa 230-0045, Japan (Shunichi Kosugi). Japan International Research Center for Agricultural Sciences, Tsukuba, Ibaraki 305-8686, Japan (Kazuhiro Sasaki). Corresponding author: Kenta Shirasawa.

## Abstract

Wild plants are often tolerant to biotic and abiotic stresses in their natural environments, whereas domesticated plants such as crops frequently lack such resilience. This difference is thought to be due to the high levels of genome heterozygosity in wild plant populations and the low levels of heterozygosity in domesticated crop species. In this study, common vetch (*Vicia sativa*) was used as a model to examine this hypothesis. The common vetch genome (2n = 14) was estimated as 1.8 Gb in size. Genome sequencing produced a reference assembly that spanned 1.5 Gb, from which 31,146 genes were predicted. Using this sequence as a reference, 24,118 single nucleotide polymorphisms were discovered in 1,243 plants from 12 natural common vetch populations in Japan. Common vetch genomes exhibited high heterozygosity at the population level, with lower levels of heterozygosity observed at specific genome regions. Such patterns of heterozygosity are thought to be essential for adaptation to different environments. These findings suggest that high heterozygosity at the population level would be required for wild plants to survive under natural conditions while allowing important gene loci to be fixed to adapt the conditions. The resources generated in this study will provide insights into *de novo* domestication of wild plants and agricultural enhancement.

**Highlight:** Sequence analysis of the common vetch *(Vicia sativa)* genome and SNP genotyping across natural populations revealed nucleotide diversity levels associated with native population environments.

## Introduction

Wild plants, including weeds that have not yet been domesticated or cultivated, generally possess characteristics that allow them to survive and propagate in their natural environments when challenged by local biotic and abiotic stresses (Mammadov *et al*., 2018). The resilience exhibited by wild plants is thought to be due to their high levels of genetic heterogeneity (Cane □ado, 2011). Indeed, genetic heterogeneity was effective in suppressing disease when populations of genetically diversified crops were planted together in the same fields (Zhu *et al*., 2000).

In contrast with wild plants, crop plants have lost their natural survival traits as a result of the extremely low levels of genetic heterogeneity found in monoculture crop species (Mundt, 2002). Therefore, disease-, insect-, and weed-controls are essential in commercial crop cultivation to reduce losses and maximize yields. This requires additional crop management costs for farmers, for example, for labor and agrochemicals. There are two main reasons for the low genetic heterogeneity in crop species. One reason is crop domestication (Izawa *et al*., 2009), in which only a few plants possessing desirable phenotypes, such as large fruit size, non-seed shattering, and long-seed dormancy, are selected from the broad genetic pools of wild plants. The second reason is selective breeding for desirable traits. While valuable for stabilizing crop phenotypes such as yield, these selective processes have reduced genetic diversity in monoculture crops by purging diverse germplasms (Fu, 2015). During domestication and selective breeding, small numbers of alleles that have large effects on phenotypic variations have often been targeted, further reducing the genetic diversity within cultivated varieties (Fernie and Yan, 2019).

While remaining more diverse than crop species, wild plant populations have also experienced loss of genetic heterogeneity at some loci, though in wild plants this is due to directional selection and genetic drift. For example, natural populations of Arabidopsis have lost genetic heterogeneity at flowering loci to synchronize flowering time (Mendez-Vigo *et al*., 2011), which is beneficial for propagation under natural conditions. This suggests that genome-wide genetic heterogeneity is not necessarily required for wild plant populations and that small numbers of loci could become fixed under certain selective conditions. This suggests that it would be possible to generate new plant populations with a) fixed domestication loci with suitable alleles for agricultural traits and b) high general levels of genetic diversity elsewhere in the genome. Such plant populations could be used as crop species, as proposed by Litrico and Violle (Litrico and Violle, 2015), and would possess natural resistance and suppression traits, as a result of high heterogeneity, that would enhance population resilience to biotic and abiotic stresses. As favorable agricultural alleles would be fixed, the benefits of genetic heterogeneity would exist alongside desirable agricultural traits. Mixtures of heterozygous plant populations have already been used as crops in allogamous species such as onion and clover. However, the potential benefits of genetic heterogeneity for autogamous plants such as legumes remain unclear.

Common vetch *(Vicia sativa),* a wild legume commonly found in open fields, was partially domesticated and cultivated in the past (Bryant and Hughes, 2011). Common vetch therefore has crop potential and can serve as a model for examination of genetic heterogeneity and domestication. The first step is to evaluate the levels of genetic heterogeneity in wild common vetch populations. However, no genome sequence data is available in common vetch. At least three different chromosome numbers (2n = 10, 12, and 14) have been reported (Ladizinsky, 1998; Ladizinsky and Waines, 1982). In this study, a reference sequence for common vetch was developed and single nucleotide polymorphism (SNP) analysis with double-digest restriction-site associated DNA sequencing (ddRAD-Seq) was used to evaluate heterogeneity in genomes of common vetch populations.

## Materials and methods

### Plant materials

A standard inbred line of common vetch *(V. sativa),* KSR5, was established from a wild plant collected from Kisarazu, Chiba, Japan, by self-pollination for more than three generations. KSR5 was used for genome and transcriptome sequencing analysis. For genetic diversity analysis, 1,243 plants were collected from 12 locations across the latitude from 31.3°N to 38.8°N in Japan (Figure 1, Supplementary Table S1). In addition, eight accessions from France, Germany, Greece, Iran, Italy, and Tunisia were obtained from the NIAS Genebank, Tsukuba, Japan (Supplementary Table S1). Genomic DNA was extracted from young leaves with a DNeasy Plant Mini Kit (Qiagen, Hilden, Germany).

**Figure 1.**
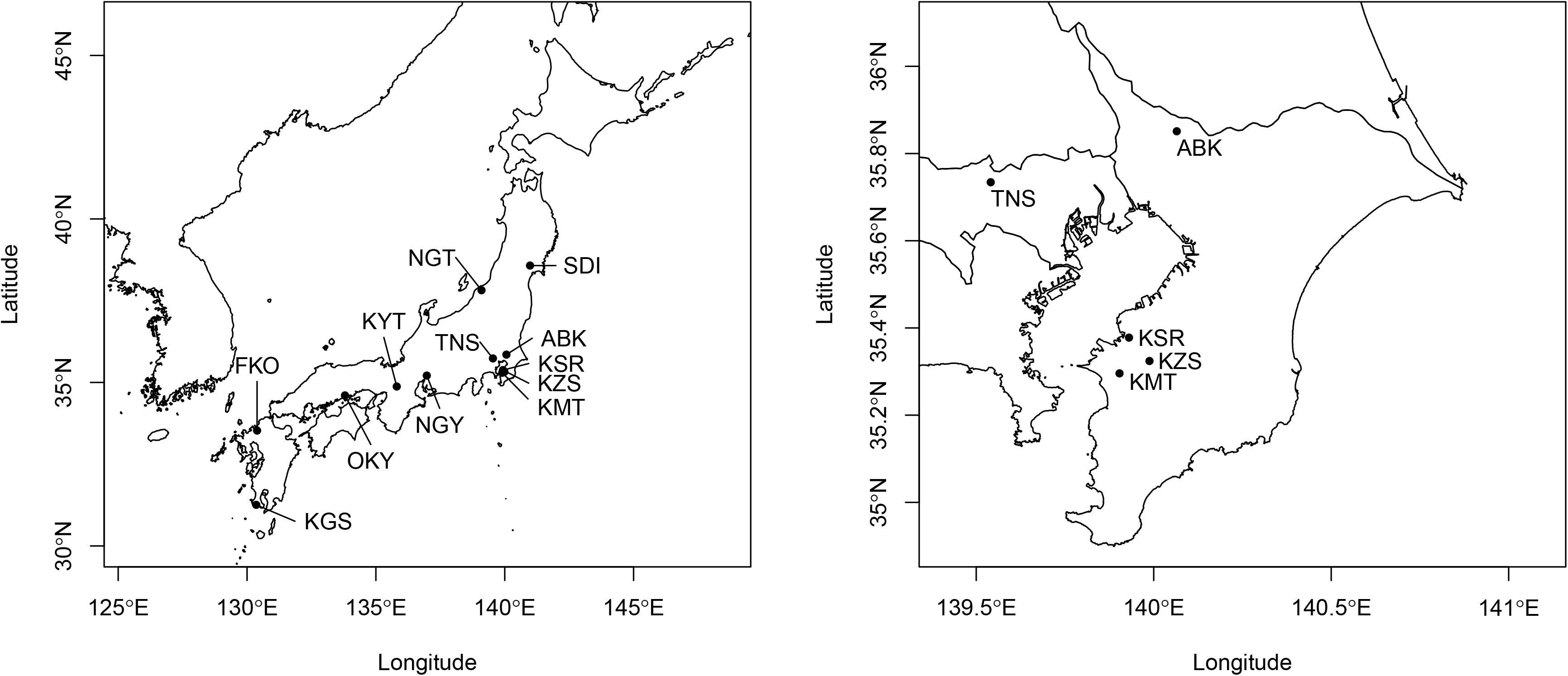
Sampling locations in Japan. Three-letter codes indicate sampling locations in Japan: ABK: Abiko, Chiba; FKO: Fukuoka; KGS: Kagoshima; KMT: Kimitsu, Chiba; KSR: Kisarazu, Chiba; KYT: Kyoto; KZS: Kazusa, Chiba; NGT: Niigata; NGY: Nagoya, Aichi; OKY: Okayama; SDI: Sendai, Miyagi; and TNS: Tanashi, Tokyo.

### Chromosome observation

Root tips of two-day-old seedlings of KSR5 were treated with 0.05% colchicine for 18 hours, fixed with 1:3 acetate:ethanol for 2 hours, and washed three times with water. Cell walls of the root tips were digested with 2% cellulase (SERVA Electrophoresis GmbH, Heidelberg, Germany), 2% macerozyme (SERVA Electrophoresis GmbH), and 0.1 M sodium acetate for four hours at 37°C. The root tip cells spread on a slide glass were fixed again with 1:3 acetate:ethanol and dried at room temperature. Chromosomes were stained with 1 ug/mL DAPI (4,6-Diamidino-2-phenylindole) in Fluoro-KEEPER Antifade Reagent (Nacalai Tesque, Kyoto, Japan) and were observed under a confocal laser scanning microscope, LSM700 (Carl Zeiss, Oberkochen, Germany). Chromosome length was measured with ImageJ (Schneider *et al*., 2012).

### Sequencing analysis of the common vetch genome

Genomic DNA from KSR5 was used to construct one paired-end (insert size of 500 bp) and four mate-pair sequencing libraries (insert sizes of 2, 5, 10, and 15 kb) in accordance with manufacturer protocols (Illumina, San Diego, CA, USA). Libraries were then sequenced using a HiSeq2000 instrument (Illumina). A long insert library for KSR5 was also prepared and sequenced on an RSII instrument (PacBio, Menlo Park, CA, USA). The paired-end sequence reads were used for genomic size estimation based on *k*-mer frequency *(k* = 17) using Jellyfish (Marcais and Kingsford, 2011). The paired-end and mate-pair reads were assembled and scaffolded with SOAPdenovo2 (Luo *et al*., 2012). Gaps, represented by Ns in the scaffold sequences, were filled by PBjelly (English *et al*., 2012) with PacBio reads, in which sequence errors were corrected with the paired-end reads by proovread (Hackl *et al*., 2014). Contaminated sequences were removed by BLASTN search (Altschul *et al*., 1990), with an E-value cutoff of 1E-10 and length coverage of ≥10%, against sequences from potential contaminating resources such as organelles (the plastid and mitochondrion genome sequences of *L. japonicus* and *V. faba:* KF042344, AP002983, JN872551, and KC189947), bacteria and fungi (NCBI bacteria and fungi), human (hg19), and artificial sequences (UniVec and PhiX). The resulting sequences that were ≥1,000 bp in size were selected and designated VSA_r1.0 as a draft common vetch genome. Completeness of the assembly was assessed with sets of a Benchmarking Universal Single-Copy Orthologs (BUSCO) (Simao *et al*., 2015).

### RNA sequencing and assembly

Total RNA was extracted from ten tissue samples (roots, seedlings, stems, apical buds, immature and mature leaves, tendrils, flower buds, flowers, and pods) using an RNeasy Mini Kit (Qiagen) and treated with RQ1 RNase-Free DNase (Promega, Madison, WI, USA) to remove contaminating genomic DNA. RNA libraries were constructed in accordance with the TruSeq Stranded mRNA Sample Preparation Guide (Illumina). Nucleotide sequences were obtained with a MiSeq instrument (Illumina) in the paired-end 301 bp mode. Low-quality reads were removed using PRINSEQ (Schmieder and Edwards, 2011) and adapter sequences were trimmed with fastx_clipper (parameter, -a AGATCGGAAGAGC) in the FASTX-Toolkit (http://hannonlab.cshl.edu/fastx_toolkit). The resulting reads were assembled using Trinity (Grabherr *et al*., 2011) with parameters of -min_contig_length 100, -group_pairs_distance 400, and -SS_lib_type RF to generate a non-redundant gene sequence set.

### Repetitive sequence and RNA coding gene analysis

A *de novo* repeat sequence database for VSA_r1.0 was built using RepeatScout (Price *et al*., 2005) (version 1.0.5). Repetitive sequences in VSA_r1.0 were searched for using RepeatMasker (version 4.0.3) (http://www.repeatmasker.org) based on known repetitive sequences registered in Repbase (Bao *et al*., 2015) and the *de novo* repeat libraries. Transfer RNA genes were predicted using tRNAscan-SE (version 1.23) (Chan and Lowe, 2019) with the default parameters, and ribosomal RNA (rRNA) genes were predicted using BLASTN searches with an E-value cutoff of 1E-10, with the *Arabidopsis thaliana* 18S rRNA (accession number: X16077) and 5.8S and 25S rRNAs (accession number: X52320) used as query sequences.

### Protein-coding gene prediction and annotation

Putative protein-coding genes in VSA_r1.0 were identified with a MAKER pipeline (version 2.31.8) (Cantarel *et al*., 2008) including *ab-initio-,* evidence-, and homology-based gene prediction methods. For this prediction, the non-redundant gene sequence set generated from the RNA-Seq analysis and peptide sequences predicted in the genomes of four Fabaceae members, namely, *Arachis duranensis* (V14167.a1.M1) (Bertioli *et al*., 2016), *Lotus japonicus* (rel. 3.0) (Sato *et al*., 2008),*Medicago truncatula* (4.0v1) (Young *et al*., 2011), and *Phaseolus vulgaris* (v1.0) (Schmutz *et al*., 2014), were used as a training data set. In addition, BRAKER1 (version 1.3) (Hoff *et al*., 2016) was used to complete the gene set for VSA_r1.0. Genes related to transposable elements (TEs) were detected using BLASTP searches against the NCBI non-redundant (nr) protein database with an E-value cutoff of 1E-10 and by using InterProScan (version 4.8) (Jones *et al*., 2014) searches against the InterPro database with an E-value cutoff of 1.0.

Putative VSA_r1.0 genes were clustered using CD-hit (version 4.6.1) (Li and Godzik, 2006) with the UniGene set of the four Fabaceae members as above with the parameters c□ = Ū0.6 and aL = 0.4. The predicted genes were annotated with plant gene ontology (GO) slim categories and euKaryotic clusters of Orthologous Groups (KOG) categories (Tatusov *et al*., 2003), and mapped onto the Kyoto Encyclopedia of Genes and Genomes (KEGG) reference pathways (Ogata *et al.*, 1999).

Gene expression was quantified by mapping the RNA-Seq reads onto VSA_r1.0 using HISAT2 (Kim *et al*., 2015) followed by normalization to determine fragments per kilobase of exon per million mapped fragments (FPKM) values using StringTie (Pertea *et al*., 2015) and Ballgown (Frazee *et al*., 2015) in accordance with the published protocol (Pertea *et al*., 2016).

### Genetic diversity analysis

Genome-wide sequence variations in wild vetch populations were analyzed by a double-digest restriction-site associated DNA sequencing (ddRAD-Seq) technique (Peterson *et al*., 2012). In accordance with the workflow established in our previous study (Shirasawa *et al*., 2016), genomic DNA samples from each line were digested with the restriction enzymes *PstI* and *Eco*RI to prepare ddRAD-Seq libraries, which were then sequenced on a HiSeq2000 (Illumina) instrument in paired-end 93 bp mode. Low-quality sequences were removed and adapters were trimmed using PRINSEQ (Schmieder and Edwards, 2011) and fastx_clipper in the FASTX-Toolkit (http://hannonlab.cshl.edu/fastx_toolkit), respectively. The remaining high-quality reads were mapped onto VSA_r1.0 as a reference using Bowtie2 (Langmead and Salzberg, 2012). The resultant sequence alignment-map format (SAM) files were converted to binary sequence alignment-map format (BAM) files and subjected to SNP calling using the mpileup option of SAMtools (Li *et al.*, 2009) and the view option of BCFtools. High-confidence SNPs were selected using VCFtools (Danecek *et al*., 2011) with the following criteria: (1) depth of coverage ≥5 for each line, (2) SNP quality scores of 999 for each locus, (3) minor allele frequency ≥0.05 for each locus, and (4) proportion of missing data <0.5 for each locus. The effects of SNPs on gene function were predicted using SnpEff v4.2 (Cingolani *et al*., 2012).

Nucleotide divergency (π) values and heterozygosity levels for SNP sites of each population were calculated using the site-pi and het options in VCFtools (Danecek *et al.*, 2011), respectively. Principal component analysis (PCA) was performed to determine the relationships among samples using TASSEL (Bradbury *et al*., 2007) and population structure was investigated using ADMIXTURE (Alexander *et al*., 2009). The R package WGCNA (Langfelder and Horvath, 2008) was used for SNP module detection.

## Results

### Chromosome number of a common vetch line, KSR5

A total of 14 chromosomes, including two mini chromosomes, were observed in metaphase cells of root tips of the standard inbred line, KSR5 (Figure 2, Table 1). Relative length of the chromosomes was measured in five cells and sorted by the length order. In accordance with the chromosome length, the 14 chromosomes were grouped into seven pairs (I to VII), suggesting that the genome of KSR5 was 2n = 14. The relative length of the longest chromosome (I) was 22.3% of the total length of haploid genome, followed by 21.0% (II), 18.6% (III), 16.1% (IV), 10.3% (V), 9.3% (VI), and 2.7% (VII).

**Figure 2.**
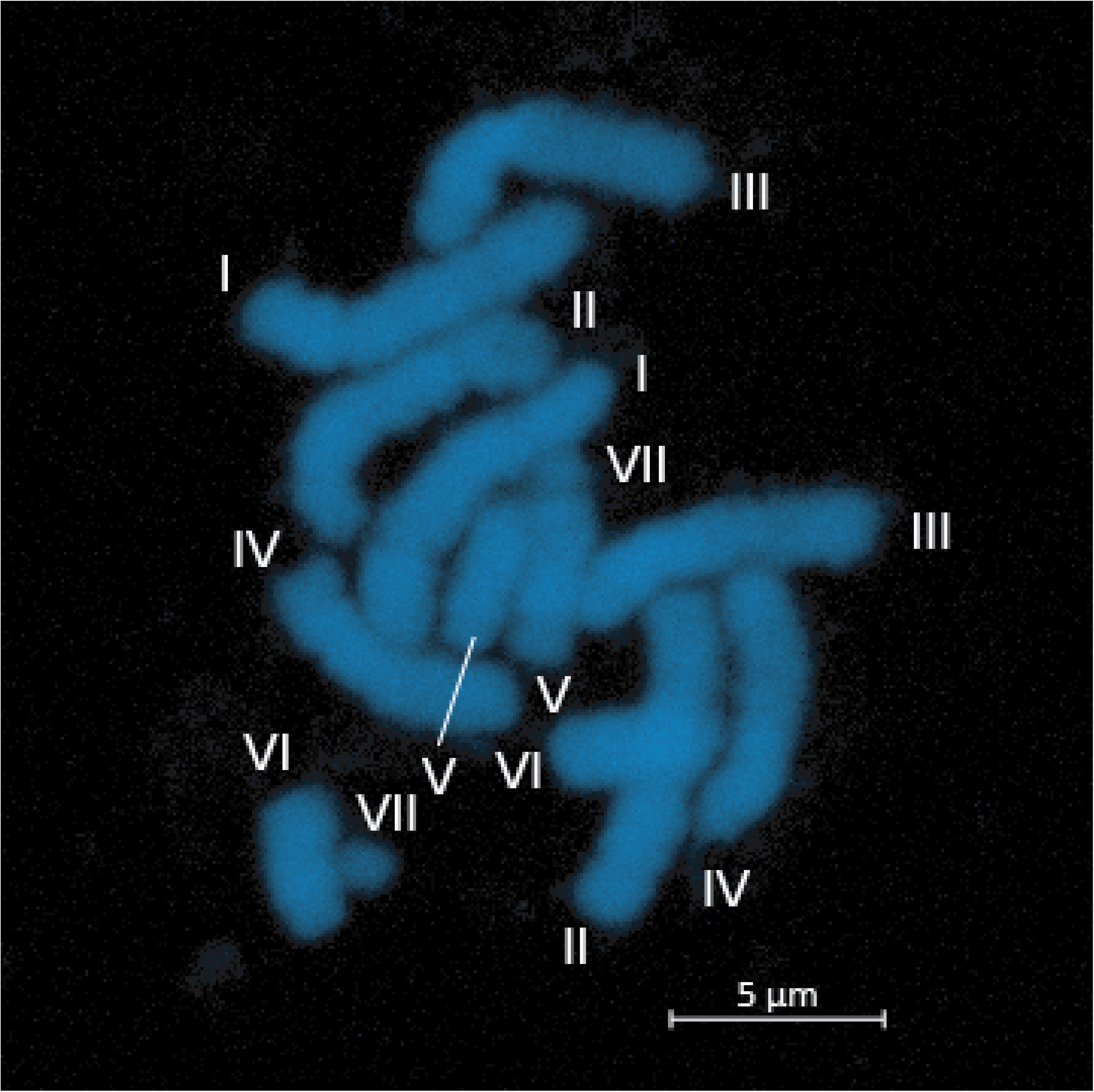
Chromosomes of the common vetch KSR5. Roman numerals indicate chromosome pairs, which order is based on chromosome length (I to VII). Bar = 5 μm.

**Table 1.** Relative chromosome length of *Vicia sativa,* KSR5

| Chromosome | Relative length (%) | S.d.* |
| --- | --- | --- |
| I | 22.3 | 0.7 |
| II | 21.0 | 0.7 |
| III | 18.6 | 1.3 |
| IV | 16.1 | 1.6 |
| V | 10.3 | 0.7 |
| VI | 9.1 | 0.6 |
| VII | 2.7 | 1.0 |

\*Standard deviation (n = 10)

### Sequencing and genome assembly

The standard inbred line of common vetch *(V. sativa),* KSR5, was sequenced. In total, 1.8 billion paired-end reads corresponding to 186.7 Gb (Supplementary Table S2) were obtained. The distribution of distinct *k*-mers (*k* = 17) showed a single main peak at multiplicities of 78 with minor peaks (Figure 3). The size of the common vetch genome was estimated to be 1,769 Mb. The paired-end reads (105× genome coverage) were assembled with mate-pair reads of four libraries (146× genome coverage in total) to obtain 6,487 thousand (k) scaffold sequences of total length 2.5 Gb with an N50 of 30.5 kb. After removing 6,421 k contaminated sequences and short scaffolds (<1 kb), sequence gaps presented by Ns in the remaining sequences were filled with PacBio long reads (3× genome coverage) to obtain a draft sequence of the common vetch genome, namely, VSA_r1.0. The total length of VSA_r1.0 was 1,541 Mb and consisted of 54,083 sequences with an N50 of 90.1 kb (Table 2). Although 513 k gaps occupied 501 Mb in total (32.5%), the gene space was well represented in accordance with BUSCO examination, indicating 94.1% ortholog completion.

**Figure 3.**
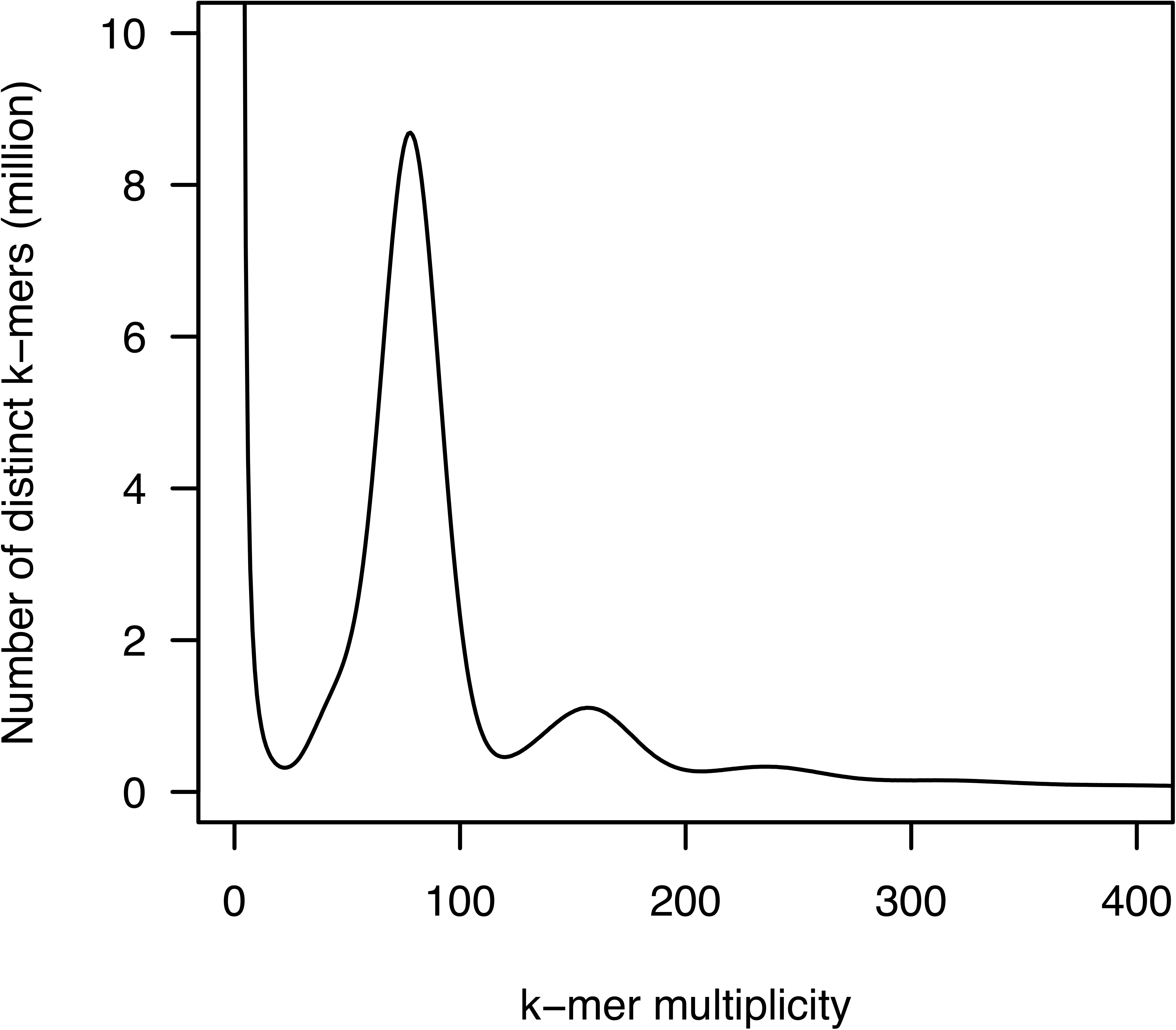
Genome size estimation for *Vicia sativa* with the distribution of the number of distinct *k*-mers (*Ä*=17) with the given multiplicity values.

**Table 2.**
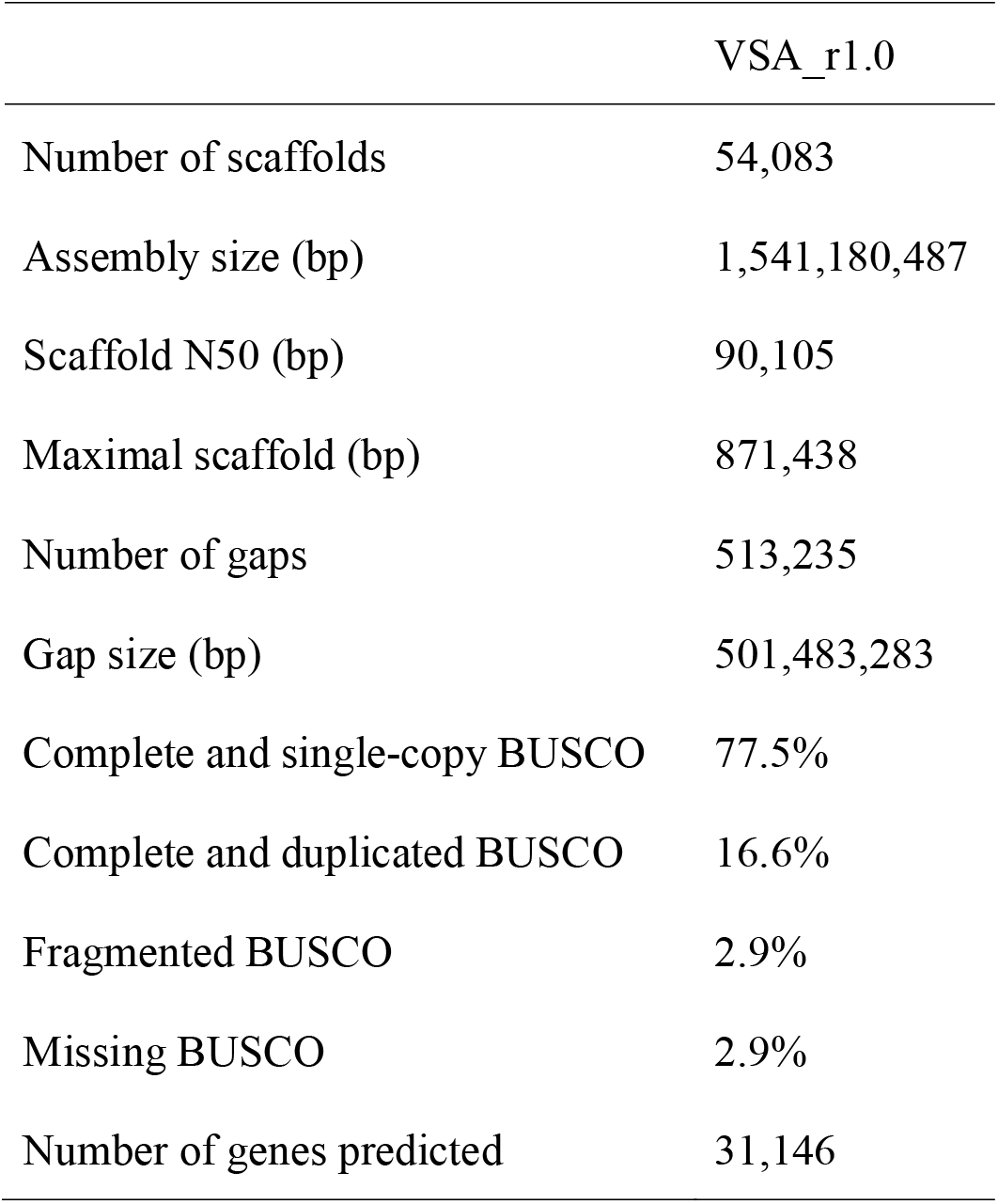
Assembly statistics of the common vetch *(Vicia sativia)* genome assembly VSA_r1.0

### Repeat sequence analysis

Sequences totaling 782 Mb (51.9%) were identified as repeat elements such as transposons and retrotransposons (Table 3). Of this, sequences totaling 267 Mb were repeat sequences reported in other organisms, and sequences in the remaining 531 Mb were uniquely identified in VSA_r1.0. Of the previously reported repeats, long terminal repeat retroelements were predominant (200 Mb). Furthermore, 109,151 simple-sequence repeats with 52,874 di-, 39,198 tri-, 12,354 tetra-, 3,414 penta-, and 1,311 hexa-nucleotide repeat motifs were also found.

**Table 3.** Repeat sequences in the VSA_r1.0 assembly

| Repeat type | Length occupied (bp) | % |
| --- | --- | --- |
| SINEs<sup>b</sup> | 85,029 | 0.0 |
| LINEs<sup>b</sup> | 10,462,622 | 0.7 |
| LTR elements<sup>b</sup> | 200,723,246 | 13.0 |
| DNA elements | 15,595,575 | 1.0 |
| Helitrons | 1,469,970 | 0.1 |
| Satellites | 17,496,670 | 1.1 |
| Simple repeats | 17,496,670 | 1.1 |
| Low complexity | 4,468,370 | 0.3 |
| Novel repeats | 531,016,543 | 34.5 |
| **Total<sup>a</sup>** | **782,834,201** | **50.8** |

<sup>a</sup>Non-redundant sequence length of the repeats overlapping in the genome.
<sup>b</sup>SINEs: short interspersed nuclear elements; LINEs: long interspersed nuclear elements; and LTR: long terminal repeat.

### Gene prediction and annotation

In total, 31,146 protein-encoding genes, with average length of 1,008 bp and N50 of 1,419 bp, were predicted in VSA_r1.0 (Table 2). For the evidence-based MAKER pipeline, 166 million (M) RNA reads from ten tissue samples (Supplementary Table S2) were assembled into 181,211 transcribed sequences and used to predict 27,880 genes (genes with .mk suffix). A further 3,266 genes were predicted using an *ab-initio*-based method (genes with .br suffix). GO classification assigned 8,878, 4,059, and 13,752 genes to the GO slim terms of biological process, cellular component, and molecular function, respectively (Supplementary Table S3). KOG analysis revealed 2,766, 4,888, and 4,424 genes with significant similarities to genes involved in information storage and processing, cellular processing and signaling, and metabolism, respectively (Supplementary Table S4). Finally, 1,720 genes were mapped to KEGG metabolic pathways (Supplementary Table S5). Gene clustering analysis revealed 5,566 gene clusters that were common to the five legume species tested *(V. sativa, A. duranensis, L. japonicus, M. truncatula,* and *P. vulgaris)* and 12,321 clusters that were unique to common vetch (Figure 4). In addition to mRNA sequences, 58 rRNA- and 1,437 tRNA-encoding genes were predicted.

**Figure 4.**
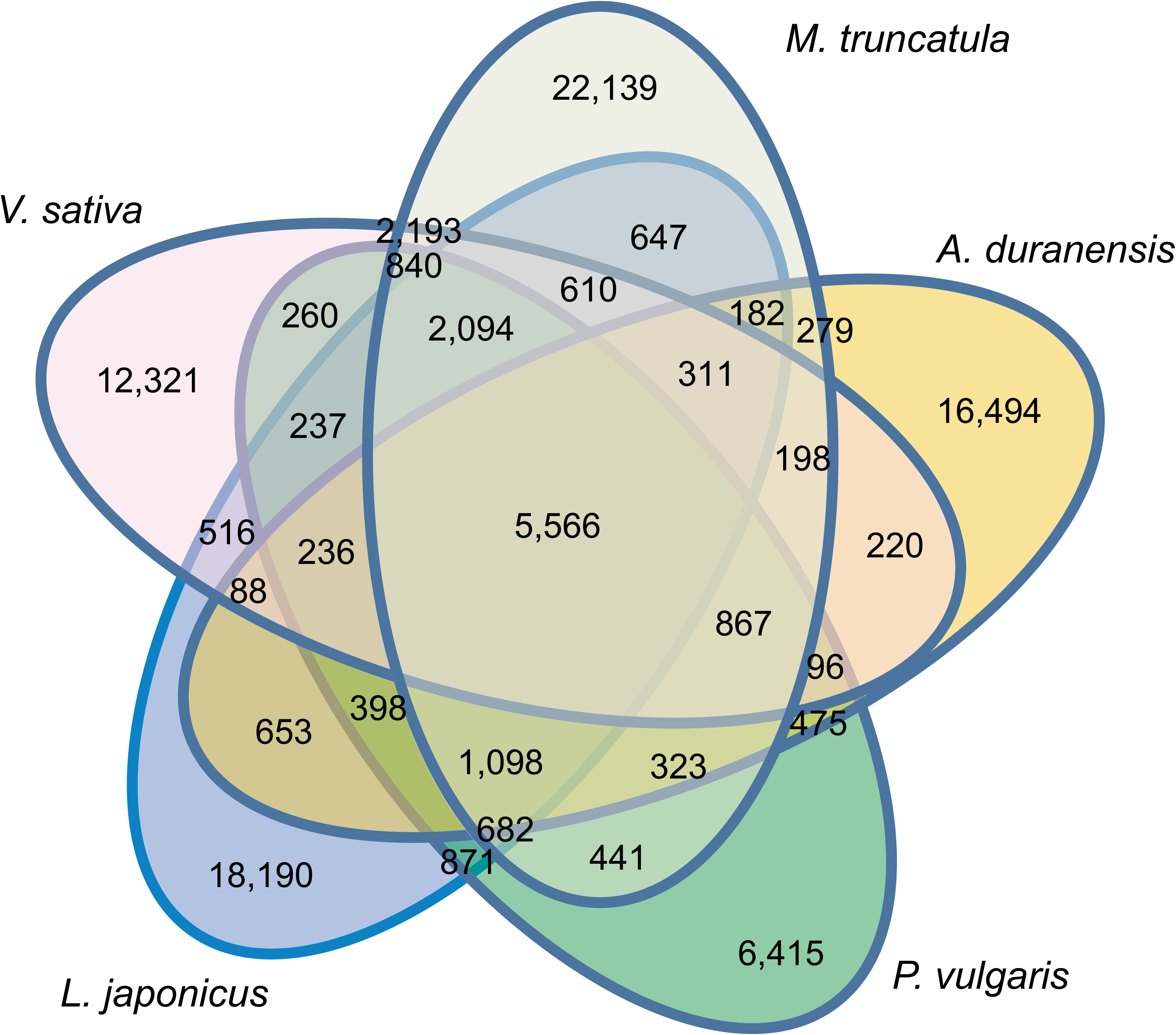
Venn diagram showing numbers of gene clusters in *Vicia sativa* and four additional Fabaceae species.

### Single nucleotide polymorphisms in natural populations

Genome-wide SNPs were identified across the 12 common vetch populations from Japan, consisting of 1,243 lines, and eight lines from France, Germany, Greece, Iran, Italy, and Tunisia from the NARO GeneBank (Tsukuba, Japan) (Supplementary Table S1). Approximately 1.1 million ddRAD-Seq reads per sample were obtained (Supplementary Table S2) and 84.4% of the reads aligned to the VSA_r1.0 reference sequence. The ddRAD-Seq reads covered 2.4 Mb (0.16%) of the reference assembly with ≥5 reads. Sequence alignments detected 46,715 high-confidence SNPs (30.9% transitions and 69.1% transversions). SNP density was calculated as 1 SNP per 51 bp. When only the 12 populations from Japan were considered, the number of SNPs decreased to 24,118 (1 SNP per 100 bp), ranging from 4,709 SNPs in the SDI population (1 SNP per 510 bp) to 10,040 SNPs in the ABK population (1 SNP per 239 bp) (Table 4).

**Table 4.**
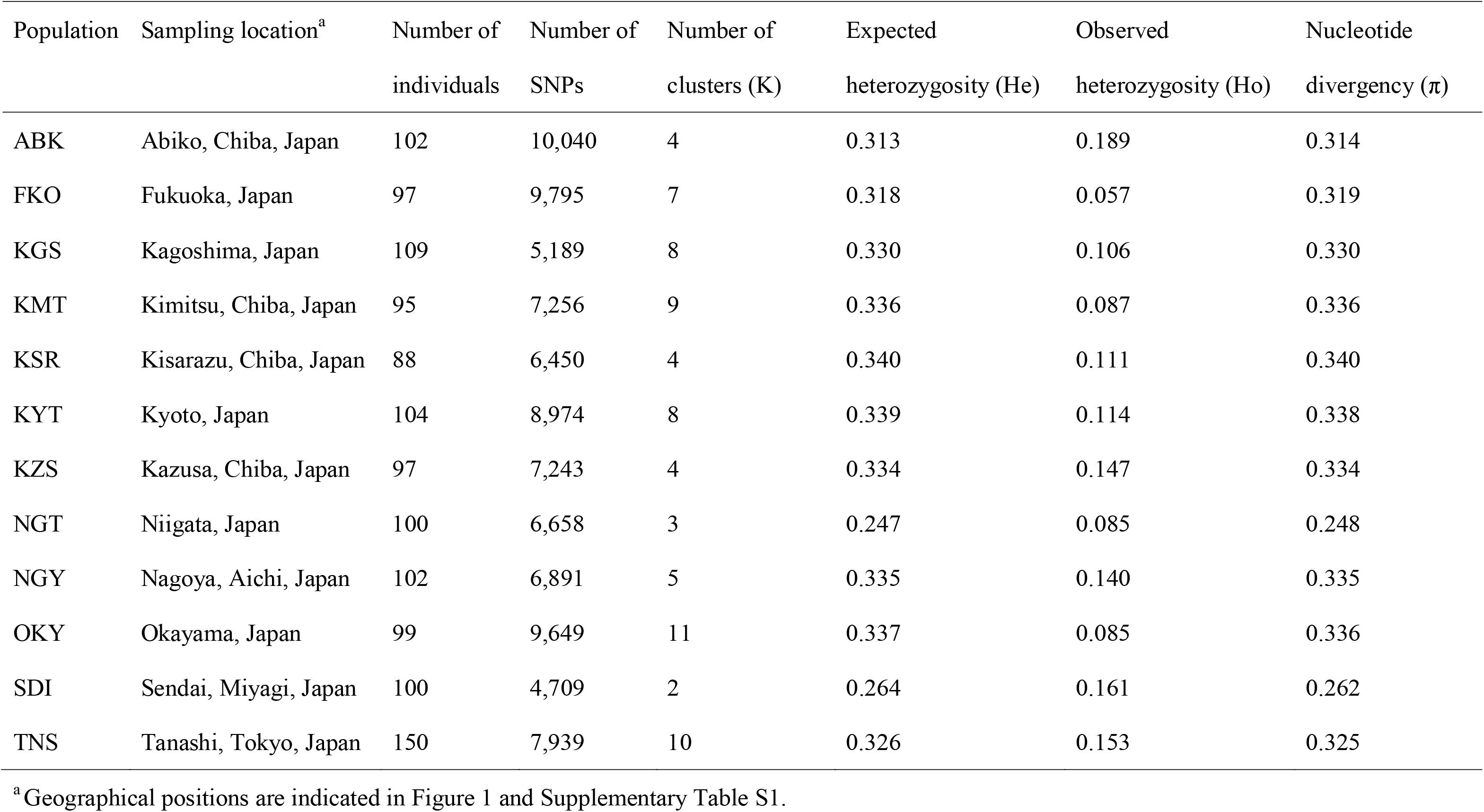
Cluster, heterozygosity, and nucleotide diversity calculated from SNPs of 12 common vetch natural populations in Japan

PCA and admixture analysis indicated that there were 2–11 subpopulations in each of the 12 populations from Japan (Figure 5, Table 3, Supplementary Figures S1). The observed heterozygosity scores were lower than the expected values (Table 4). Nucleotide divergency scores (π) at SNP sites were similarly distributed across ten of the populations from Japan, with median values of 0.31–0.34. The remaining two populations, NGT and SDI, exhibited median values of ~0.25 (Table 4). Of the 46,715 high-confidence SNPs, 24,118 clustered according to their π scores to generate 82 modules (Supplementary Figure S2). Of these, the π scores of one cluster, ‘cyan’, which contained 190 SNPs, negatively correlated with the latitude of sampling location (Figure 1 and 6). In total, 88 genes were associated with the 190 SNPs, and one of the genes (Vsa_sc30698.1_g030.1.mk) showed sequence similarity to the Arabidopsis gene for a MADS-box protein, SUPPRESSOR OF OVEREXPRESSION OF CONSTANS1 (SOC1), known to be involved in the flowering pathway in plants. Vsa_sc30698.1_g030.1.mk was predominantly transcribed in tendrils (FPKM = 5.0) followed by apical buds (0.5) and stems (0.4), whereas no expression was observed in the other seven tissues, i.e., roots, seedlings, immature and mature leaves, flower buds, flowers, and pods.

**Figure 5.**
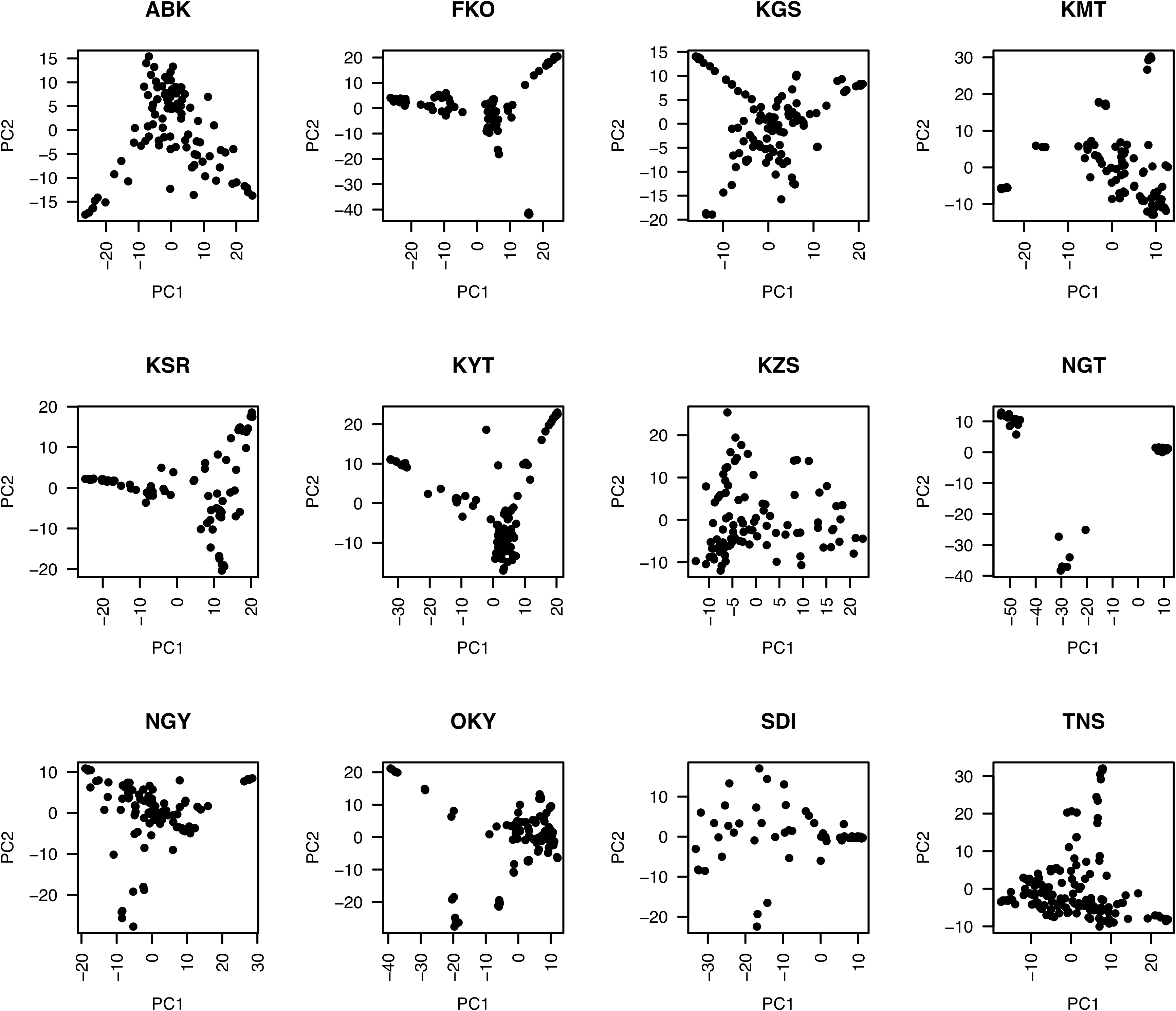
Principal component analysis of 12 natural populations of *Vicia sativa* from Japan.

**Figure 6.**
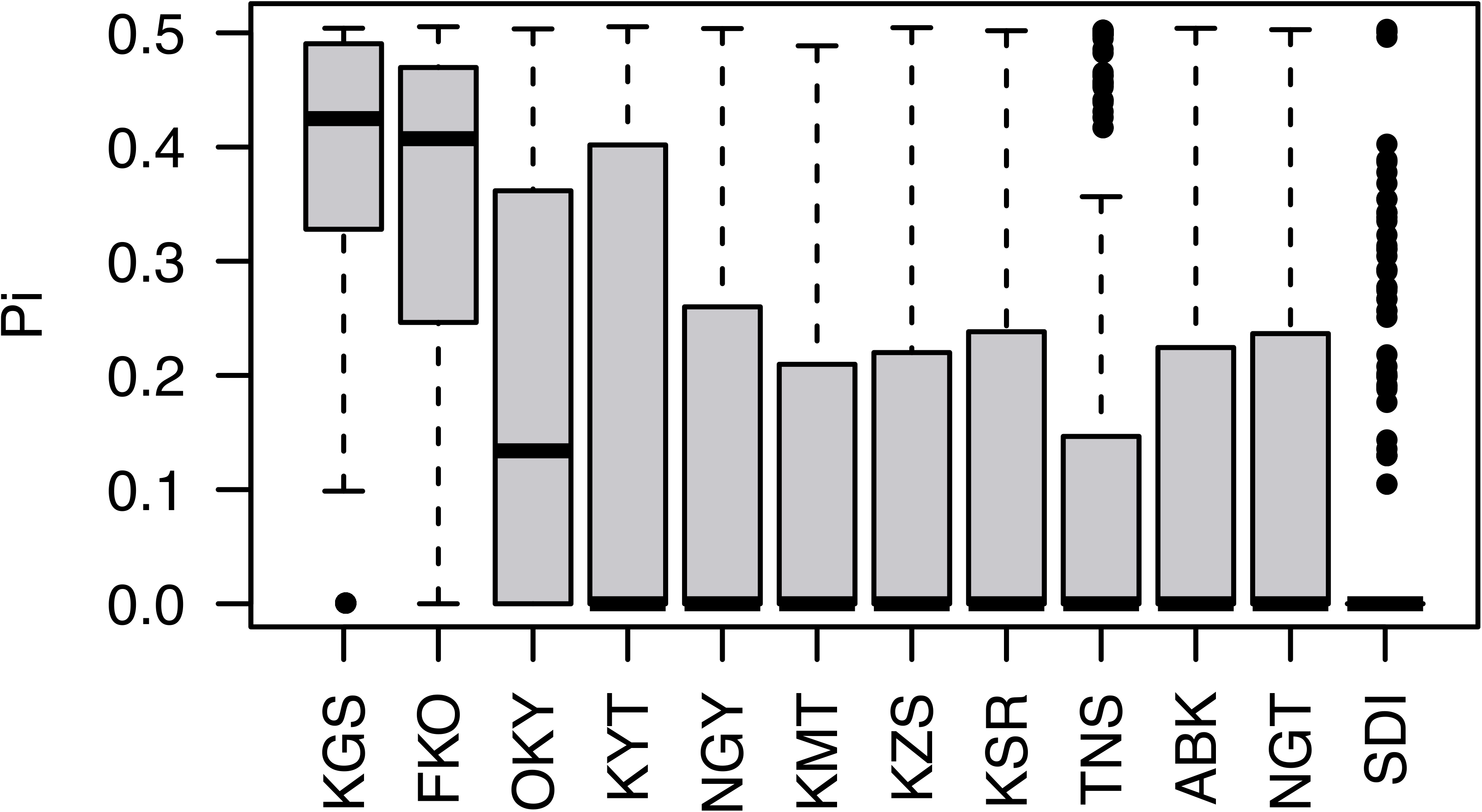
Nucleotide diversity (π) of the SNP module ‘cyan’ (n=190) across 12 natural populations of *Vicia sativa* in Japan. Three-letter codes indicate sampling locations in Japan: ABK: Abiko, Chiba; FKO: Fukuoka; KGS: Kagoshima; KMT: Kimitsu, Chiba; KSR: Kisarazu, Chiba; KYT: Kyoto; KZS: Kazusa, Chiba; NGT: Niigata; NGY: Nagoya, Aichi; OKY: Okayama; SDI: Sendai, Miyagi; and TNS: Tanashi, Tokyo.

## Discussion

A draft: common vetch *(V. sativa)* genome sequence was generated in this study. Although several legume genome sequences were released previously (Bauchet *et al*., 2019), this is the first report of a genome from the genus *Vicia,* which contains several agronomically important legume crops such as fava bean *(V. faba). Vicia* genomes are large (e.g., 1.8 Gb for *V. sativa* and 13 Gb for *V. faba)* due to their massive repetitive sequences, including TEs (Bryant and Hughes, 2011; Hill *et al.*, 2005;

Nouzova *et al*., 2001; Pearce *et al*., 1996), hampering *de novo* genome assembly in this genus (Bauchet *et al*., 2019). As might therefore be expected, more than half of the *V. sativa* genome assembly was comprised of repetitive sequences (Table 3). The assembly contained up to 54,083 contig sequences and included 513 k gaps occupying >500 Mb (Table 2). The short-read technology employed for sequencing might therefore be insufficient to span the repeats. Although construction of contiguous sequences from the short reads was challenging, a near complete gene set was successfully identified in the assembly (Table 2). Whereas it was impossible to compare the genome structure of common vetch with those of relatives due to the fragmented genome sequences, clustering analysis of the gene sequences would provide insights into the gene homoeology in legume species (Figure 4). The genome resources developed in this study will be invaluable for forthcoming gene discovery studies, such as transcriptome analysis and allele mining, in *Vicia.*

We reproducibly observed seven pairs of chromosomes (I to VII) in the root-tip cells of KSR5 (Figure 2), among of which one pair (VII) was so small occupying only 2.7% of the total length of the seven chromosome pairs (Table 1). One type of mini chromosomes, so called B chromosomes which are comprised of repetitive sequence, have been reported in numerous groups of plants so far, but the biological function has not been known (Houben, 2017). B chromosomes are not necessary for the growth and normal development of organisms and show non-Mendelian inheritance patterns (Houben, 2017). This could be one of the reasons for the different chromosome numbers in *Vicia sativa* (Ladizinsky, 1998; Ladizinsky and Waines, 1982; Navratilova *et al*., 2003). Further chromosome observations and fluorescence in situ hybridization with the repetitive sequences as probes across multiple lines would characterize and identify the mini chromosomes observed in this study. Alternatively, sterility of F1 hybrids derived from crosses between plants with different chromosome numbers should be analyzed to gain insights into the function of the small chromosomes.

Twelve common vetch populations from Japan were examined, each of which contained 2—11 subpopulations (Figure 5, Table 4, Supplementary Figures S4). This suggested that the numbers of founder plants were limited even in populations grown under natural environmental conditions. Heterozygosity is thought to contribute strongly to the survival of plant populations under natural conditions (Canc□ado, 2011). Here, the observed heterozygosity was lower than expected (Table 4), indicating that heterozygosity in common vetch populations was high at the population level but low at the individual level due to self-pollination. This suggested that high heterozygosity at the population level is sufficient to allow adaptation and survival under natural conditions in autogamous common vetch.

Human domestication of wild plant species for agriculture involved selection of individual plants with desirable traits (Izawa *et al*., 2009; Vaughan *et al*., 2007). More recently, elite cultivars have been developed with enhanced yield performance to satisfy global food requirements (Hickey *et al*., 2019). The successive selection of small numbers of individual plants during these processes produced severe bottleneck effects and resulted in decreased genetic diversity and lower tolerance to biotic and abiotic stresses (Canc□ado, 2011). Heterozygosity at specific genome regions was also lost in some wild plants (Figure 6), as reported previously (Mendez-Vigo *et al*., 2011). This suggested that genome-wide genetic heterogeneity is not necessarily required for plants to survive under natural conditions. Recent studies have proposed *de novo-,* super-, or neo-domestication (Fernie and Yan, 2019; Hickey *et al*., 2019; Vaughan *et al*., 2007), whereby genetic loci for agronomically important traits are introduced to cultivated crop varieties from wild plants. However, the high genetic heterozygosity levels from the wild donor plants should be retained during the development of new crops to avoid the bottleneck effects sustained during historic domestication of crop varieties (Litrico and Violle, 2015). Therefore, we propose that new domestication of wild plants should retain high heterozygosity at the population level to capitalize on beneficial traits that increase tolerance to abiotic and biotic stresses, but that agronomically important genetic loci should be fixed to maximize crop potential. The resources generated in this study will provide insights into the *de novo* domestication of wild plants to develop enhanced crop varieties.

## Supplementary Data

**Supplementary Table S1** Plant materials.

**Supplementary Table S2** Genome and transcriptome data.

**Supplementary Table S3** Number of KOG functions for protein-encoding genes.

**Supplementary Table S4** Number of genes mapped to KEGG pathways.

**Supplementary Table S5** Number of GO terms for protein-encoding genes.

**Supplementary Figure S1** Cross-validation errors for 12 natural populations of *Vicia sativa* from Japan in admixture analysis.

**Supplementary Figure S2** Nucleotide diversity of SNP modules across 12 natural populations of *Vicia sativa* from Japan.

## Acknowledgments

We are grateful to Dr. H. Masumoto (Kazusa DNA Research Institute) for his kind support. We thank T. Fujishiro, K. Kawashima, Y. Kishida, M. Kohara, C. Minami, S. Nakayama, K. Nanri, S. Sasamoto, C. Takahashi, H. Tsuruoka, A. Watanabe, and M. Yamada (Kazusa DNA Research Institute) and staff of the Department of Technical Development at the Institute for Sustainable Agro-ecosystem Services (University of Tokyo) for their technical assistance. Plant materials were provided by the NIAS GeneBank (Tsukuba, Japan). This work was supported by KAKENHI (24710237 and 221S0002) and the Kazusa DNA Research Institute Foundation.

## Data Availability

Sequence data are available from the Sequence Read Archive (DRA) of DNA Data Bank of Japan (DDBJ) under accession numbers DRA004347 for whole genome sequencing, DRA004313 for RNA-Seq, and DRA004301-DRA004312 for ddRAD-Seq (Supplementary Table S2). The DDBJ accession numbers of the assembled sequences are BLWO01000001-BLWO01054083. Genome information is available at Plant GARDEN (https://plantgarden.jp).

## References

Alexander DH, Novembre J, Lange K. 2009. Fast model-based estimation of ancestry in unrelated individuals. Genome Res 19, 1655–1664.

Altschul SF, Gish W, Miller W, Myers EW, Lipman DJ. 1990. Basic local alignment search tool. J Mol Biol 215, 403–410.

Bao W, Kojima KK, Kohany O. 2015. Repbase Update, a database of repetitive elements in eukaryotic genomes. Mob DNA 6, 11.

Bauchet GJ, Bett KE, Cameron CT, Campbell JD, Cannon EKS, Cannon SB, Carlson JW, Chan A, Cleary A, Close TJ, Cook DR, Cooksey AM, Coyne C, Dash S, Dickstein R, Farmer AD, Fernáπdez Baca D, Hokin S, Jones ES, Kang Y, Monteros MJ, Muñoz □Amatriaín M, Mysore KS, Pislariu CI, Richards C, Shi A, Town CD, Udvardi M, Wettberg EB, Young ND, Zhao PX. 2019. The future of legume genetic data resources: Challenges, opportunities, and priorities. Legume Science 1.

Bertioli DJ, Cannon SB, Froenicke L, Huang G, Farmer AD, Cannon EK, Liu X, Gao D, Clevenger J, Dash S, Ren L, Moretzsohn MC, Shirasawa K, Huang W, Vidigal B, Abernathy B, Chu Y, Niederhuth CE, Umale P, Araujo AC, Kozik A, Kim KD, Burow MD, Varshney RK, Wang X, Zhang X, Barkley N, Guimaraes PM, Isobe S, Guo B, Liao B, Stalker HT, Schmitz RJ, Scheffler BE, Leal-Bertioli SC, Xun X, Jackson SA, Michelmore R, Ozias-Akins P. 2016. The genome sequences of Arachis duranensis and Arachis ipaensis, the diploid ancestors of cultivated peanut. Nat Genet 48, 438–446.

Bradbury PJ, Zhang Z, Kroon DE, Casstevens TM, Ramdoss Y, Buckler ES. 2007. TASSEL: software for association mapping of complex traits in diverse samples. Bioinformatics 23, 2633–2635.

Bryant JA, Hughes SG. 2011. Vicia. In: Kole C, ed. Wild Crop Relatives: Genomic and Breeding Resources. Berlin Heidelberg: Springer-Verlag, 273–289.

Canc□ado G. 2011. The Importance of Genetic Diversity to Manage Abiotic Stress. In: Shanker A, ed. Abiotic Stress in Plants - Mechanisms and Adaptations: InTech, 351–366.

Cantarel BL, Korf I, Robb SM, Parra G, Ross E, Moore B, Holt C, Sanchez Alvarado A, Yandell M. 2008. MAKER: an easy-to-use annotation pipeline designed for emerging model organism genomes. Genome Res 18, 188–196.

Chan PP, Lowe TM. 2019. tRNAscan-SE: Searching for tRNA Genes in Genomic Sequences. Methods Mol Biol 1962, 1–14.

Cingolani P, Platts A, Wang le L, Coon M, Nguyen T, Wang L, Land SJ, Lu X, Ruden DM. 2012. A program for annotating and predicting the effects of single nucleotide polymorphisms, SnpEff: SNPs in the genome of Drosophila melanogaster strain w1118; iso-2; iso-3. Fly (Austin) 6, 80–92.

Danecek P, Auton A, Abecasis G, Albers CA, Banks E, DePristo MA, Handsaker RE, Lunter G, Marth GT, Sherry ST, McVean G, Durbin R, Genomes Project Analysis G. 2011. The variant call format and VCFtools. Bioinformatics 27, 2156–2158.

English AC, Richards S, Han Y, Wang M, Vee V, Qu J, Qin X, Muzny DM, Reid JG, Worley KC, Gibbs RA. 2012. Mind the gap: upgrading genomes with Pacific Biosciences RS long-read sequencing technology. PLoS One 7, e47768.

Fernie AR, Yan J. 2019. De Novo Domestication: An Alternative Route toward New Crops for the Future. Mol Plant 12, 615–631.

Frazee AC, Pertea G, Jaffe AE, Langmead B, Salzberg SL, Leek JT. 2015. Ballgown bridges the gap between transcriptome assembly and expression analysis. Nat Biotechnol 33, 243–246.

Fu YB. 2015. Understanding crop genetic diversity under modern plant breeding. Theor Appl Genet 128, 2131–2142.

Grabherr MG, Haas BJ, Yassour M, Levin JZ, Thompson DA, Amit I, Adiconis X, Fan L, Raychowdhury R, Zeng Q, Chen Z, Mauceli E, Hacohen N, Gnirke A, Rhind N, di Palma F, Birren BW, Nusbaum C, Lindblad-Toh K, Friedman N, Regev A. 2011. Full-length transcriptome assembly from RNA-Seq data without a reference genome. Nat Biotechnol 29, 644–652.

Hackl T, Hedrich R, Schultz J, Forster F. 2014. proovread: large-scale high-accuracy PacBio correction through iterative short read consensus. Bioinformatics 30, 3004–3011.

Hickey LT, A Nh, Robinson H, Jackson SA, Leal-Bertioli SCM, Tester M, Gao C, Godwin ID, Hayes BJ, Wulff BBH. 2019. Breeding crops to feed 10 billion. Nat Biotechnol 37, 744–754.

Hill P, Burford D, Martin DM, Flavell AJ. 2005. Retrotransposon populations of Vicia species with varying genome size. Mol Genet Genomics 273, 371–381.

Hoff KJ, Lange S, Lomsadze A, Borodovsky M, Stanke M. 2016. BRAKER1: Unsupervised RNA-Seq-Based Genome Annotation with GeneMark-ET and AUGUSTUS. Bioinformatics 32, 767–769.

Houben A. 2017. B Chromosomes - A Matter of Chromosome Drive. Front Plant Sci 8, 210.

Izawa T, Konishi S, Shomura A, Yano M. 2009. DNA changes tell us about rice domestication. Curr Opin Plant Biol 12, 185–192.

Jones P, Binns D, Chang HY, Fraser M, Li W, McAnulla C, McWilliam H, Maslen J, Mitchell A, Nuka G, Pesseat S, Quinn AF, Sangrador-Vegas A, Scheremetjew M, Yong SY, Lopez R, Hunter S. 2014. InterProScan 5: genome-scale protein function classification. Bioinformatics 30, 1236–1240.

Kim D, Langmead B, Salzberg SL. 2015. HISAT: a fast spliced aligner with low memory requirements. Nat Methods 12, 357–360.

Ladizinsky G. 1998. Plant Evolution under Domestication. Netherlands: Kluwer Academic Publishers.

Ladizinsky G, Waines G. 1982. Seed protein polymorphism inVicia sativa agg. (Fabaceae). Plant Systematics and Evolution 141, 1–5.

Langfelder P, Horvath S. 2008. WGCNA: an R package for weighted correlation network analysis. BMC Bioinformatics 9, 559.

Langmead B, Salzberg SL. 2012. Fast gapped-read alignment with Bowtie 2. Nat Methods 9, 357–359.

Li H, Handsaker B, Wysoker A, Fennell T, Ruan J, Homer N, Marth G, Abecasis G, Durbin R, Genome Project Data Processing S. 2009. The Sequence Alignment/Map format and SAMtools. Bioinformatics 25, 2078–2079.

Li W, Godzik A. 2006. Cd-hit: a fast program for clustering and comparing large sets of protein or nucleotide sequences. Bioinformatics 22, 1658–1659.

Litrico I, Violle C. 2015. Diversity in Plant Breeding: A New Conceptual Framework. Trends Plant Sci 20, 604–613.

Luo R, Liu B, Xie Y, Li Z, Huang W, Yuan J, He G, Chen Y, Pan Q, Liu Y, Tang J, Wu G, Zhang H, Shi Y, Liu Y, Yu C, Wang B, Lu Y, Han C, Cheung DW, Yiu SM, Peng S, Xiaoqian Z, Liu G, Liao X, Li Y, Yang H, Wang J, Lam TW, Wang J. 2012. SOAPdenovo2: an empirically improved memory-efficient short-read de novo assembler. Gigascience 1, 18.

Mammadov J, Buyyarapu R, Guttikonda SK, Parliament K, Abdurakhmonov IY, Kumpatla SP. 2018. Wild Relatives of Maize, Rice, Cotton, and Soybean: Treasure Troves for Tolerance to Biotic and Abiotic Stresses. Front Plant Sci 9, 886.

Marcais G, Kingsford C. 2011. A fast, lock-free approach for efficient parallel counting of occurrences of k-mers. Bioinformatics 27, 764–770.

Mendez-Vigo B, Pico FX, Ramiro M, Martinez-Zapater JM, Alonso-Blanco C. 2011. Altitudinal and climatic adaptation is mediated by flowering traits and FRI, FLC, and PHYC genes in Arabidopsis. Plant Physiol 157, 1942–1955.

Mundt CC. 2002. Use of multiline cultivars and cultivar mixtures for disease management. Annu Rev Phytopathol 40, 381–410.

Navratilova A, Neumann P, Macas J. 2003. Karyotype analysis of four Vicia species using in situ hybridization with repetitive sequences. Ann Bot 91, 921–926.

Nouzova M, Neumann P, Navratilova A, Galbraith DW, Macas J. 2001. Microarray-based survey of repetitive genomic sequences in Vicia spp. Plant Mol Biol 45, 229–244.

Ogata H, Goto S, Sato K, Fujibuchi W, Bono H, Kanehisa M. 1999. KEGG: Kyoto Encyclopedia of Genes and Genomes. Nucleic Acids Res 27, 29–34.

Pearce SR, Harrison G, Li D, Heslop-Harrison J, Kumar A, Flavell AJ. 1996. The Ty1-copia group retrotransposons in Vicia species: copy number, sequence heterogeneity and chromosomal localisation. Mol Gen Genet 250, 305–315.

Pertea M, Kim D, Pertea GM, Leek JT, Salzberg SL. 2016. Transcript-level expression analysis of RNA-seq experiments with HISAT, StringTie and Ballgown. Nat Protoc 11, 1650–1667.

Pertea M, Pertea GM, Antonescu CM, Chang TC, Mendell JT, Salzberg SL. 2015. StringTie enables improved reconstruction of a transcriptome from RNA-seq reads. Nat Biotechnol 33, 290–295.

Peterson BK, Weber JN, Kay EH, Fisher HS, Hoekstra HE. 2012. Double digest RADseq: an inexpensive method for de novo SNP discovery and genotyping in model and non-model species. PLoS One 7, e37135.

Price AL, Jones NC, Pevzner PA. 2005. De novo identification of repeat families in large genomes. Bioinformatics 21 Suppl 1, i351–358.

Sato S, Nakamura Y, Kaneko T, Asamizu E, Kato T, Nakao M, Sasamoto S, Watanabe A, Ono A, Kawashima K, Fujishiro T, Katoh M, Kohara M, Kishida Y, Minami C, Nakayama S, Nakazaki N, Shimizu Y, Shinpo S, Takahashi C, Wada T, Yamada M, Ohmido N, Hayashi M, Fukui K, Baba T, Nakamichi T, Mori H, Tabata S. 2008. Genome structure of the legume, Lotus japonicus. DNA Res 15, 227–239.

Schmieder R, Edwards R. 2011. Quality control and preprocessing of metagenomic datasets. Bioinformatics 27, 863–864.

Schmutz J, McClean PE, Mamidi S, Wu GA, Cannon SB, Grimwood J, Jenkins J, Shu S, Song Q, Chavarro C, Torres-Torres M, Geffroy V, Moghaddam SM, Gao D, Abernathy B, Barry K, Blair M, Brick MA, Chovatia M, Gepts P, Goodstein DM, Gonzales M, Hellsten U, Hyten DL, Jia G, Kelly JD, Kudrna D, Lee R, Richard MM, Miklas PN, Osorno JM, Rodrigues J, Thareau V, Urrea CA, Wang M, Yu Y, Zhang M, Wing RA, Cregan PB, Rokhsar DS, Jackson SA. 2014. A reference genome for common bean and genome-wide analysis of dual domestications. Nat Genet 46, 707–713.

Schneider CA, Rasband WS, Eliceiri KW. 2012. NIH Image to ImageJ: 25 years of image analysis. Nat Methods 9, 671–675.

Shirasawa K, Hirakawa H, Isobe S. 2016. Analytical workflow of double-digest restriction site-associated DNA sequencing based on empirical and in silico optimization in tomato. DNA Res 23, 145–153.

Simao FA, Waterhouse RM, Ioannidis P, Kriventseva EV, Zdobnov EM. 2015. BUSCO: assessing genome assembly and annotation completeness with single-copy orthologs. Bioinformatics 31, 3210–3212.

Tatusov RL, Fedorova ND, Jackson JD, Jacobs AR, Kiryutin B, Koonin EV, Krylov DM, Mazumder R, Mekhedov SL, Nikolskaya AN, Rao BS, Smirnov S, Sverdlov AV, Vasudevan S, Wolf YI, Yin JJ, Natale DA. 2003. The COG database: an updated version includes eukaryotes. BMC Bioinformatics 4, 41.

Vaughan DA, Balazs E, Heslop-Harrison JS. 2007. From crop domestication to super-domestication. Ann Bot 100, 893–901.

Young ND, Debelle F, Oldroyd GE, Geurts R, Cannon SB, Udvardi MK, Benedito VA, Mayer KF, Gouzy J, Schoof H, Van de Peer Y, Proost S, Cook DR, Meyers BC, Spannagl M, Cheung F, De Mita S, Krishnakumar V, Gundlach H, Zhou S, Mudge J, Bharti AK, Murray JD, Naoumkina MA, Rosen B, Silverstein KA, Tang H, Rombauts S, Zhao PX, Zhou P, Barbe V, Bardou P, Bechner M, Bellec A, Berger A, Berges H, Bidwell S, Bisseling T, Choisne N, Couloux A, Denny R, Deshpande S, Dai X, Doyle JJ, Dudez AM, Farmer AD, Fouteau S, Franken C, Gibelin C, Gish J, Goldstein S, Gonzalez AJ, Green PJ, Hallab A, Hartog M, Hua A, Humphray SJ, Jeong DH, Jing Y, Jocker A, Kenton SM, Kim DJ, Klee K, Lai H, Lang C, Lin S, Macmil SL, Magdelenat G, Matthews L, McCorrison J, Monaghan EL, Mun JH, Najar FZ, Nicholson C, Noirot C, O’Bleness M, Paule CR, Poulain J, Prion F, Qin B, Qu C, Retzel EF, Riddle C, Sallet E, Samain S, Samson N, Sanders I, Saurat O, Scarpelli C, Schiex T, Segurens B, Severin AJ, Sherrier DJ, Shi R, Sims S, Singer SR, Sinharoy S, Sterck L, Viollet A, Wang BB, Wang K, Wang M, Wang X, Warfsmann J, Weissenbach J, White DD, White JD, Wiley GB, Wincker P, Xing Y, Yang L, Yao Z, Ying F, Zhai J, Zhou L, Zuber A, Denarie J, Dixon RA, May GD, Schwartz DC, Rogers J, Quetier F, Town CD, Roe BA. 2011. The Medicago genome provides insight into the evolution of rhizobial symbioses. Nature 480, 520–524.

Zhu Y, Chen H, Fan J, Wang Y, Li Y, Chen J, Fan J, Yang S, Hu L, Leung H, Mew TW, Teng PS, Wang Z, Mundt CC. 2000. Genetic diversity and disease control in rice. Nature 406, 718–722.

