## Supplementary Figure for "Genome features of common vetch (*Vicia sativa*) in natural habitats"

[†]Present address: RIKEN, Yokohama, Kanagawa 230-0045, Japan

**Supplementary Table S1** Plant materials
**Supplementary Table S2** Genome and transcriptome data
**Supplementary Table S3** Number of KOG functions for protein-encoding genes
**Supplementary Table S4** Number of genes mapped to KEGG pathways
**Supplementary Table S5** Number of GO terms for protein-encoding genes

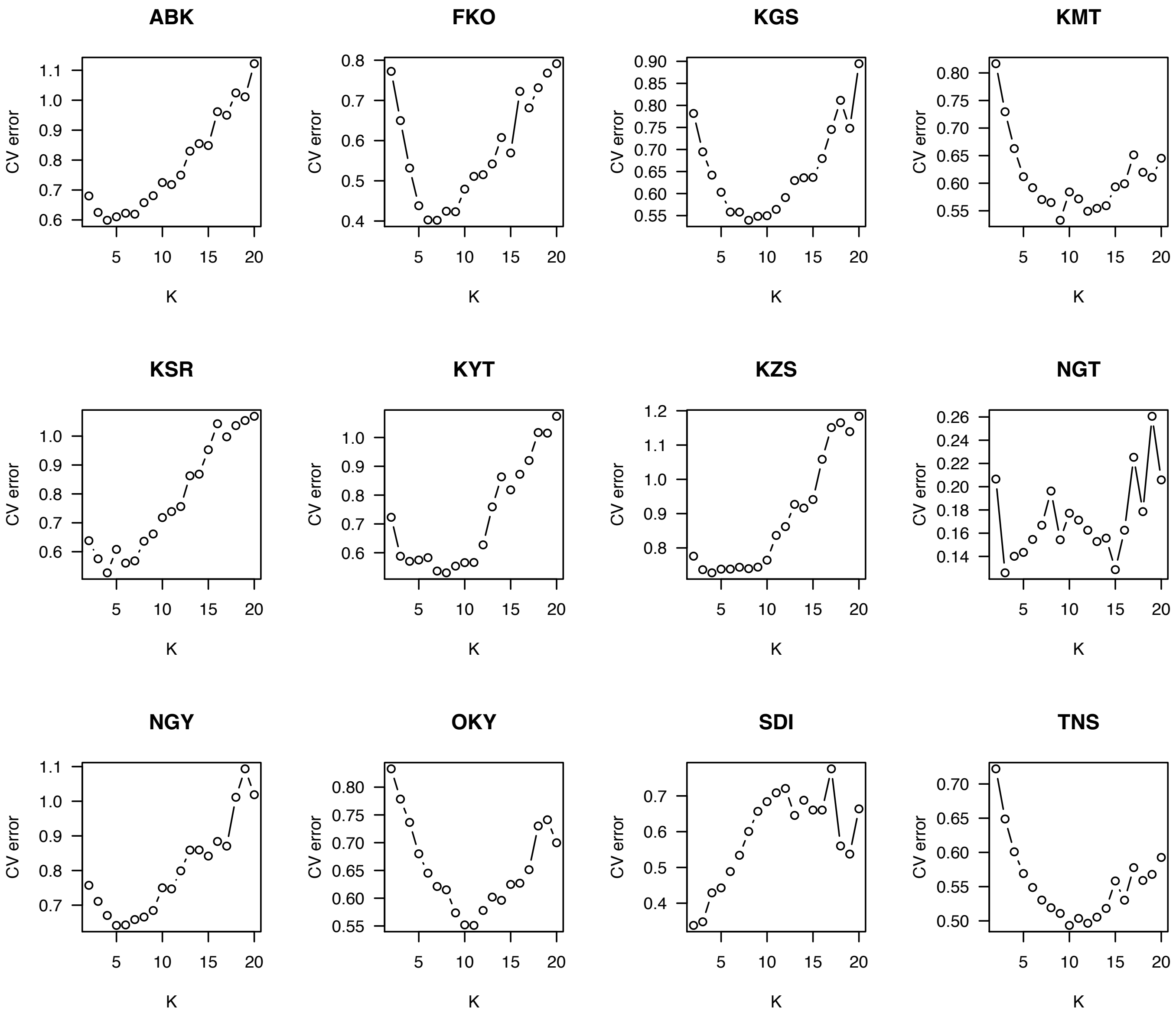


**Supplementary Figure S1** Cross-validation errors for 12 natural populations of *Vicia sativa* from Japan in admixture analysis.

Three-letter codes indicate sampling location in Japan as shown in Supplementary Table S1 and Supplementary Figure S1.

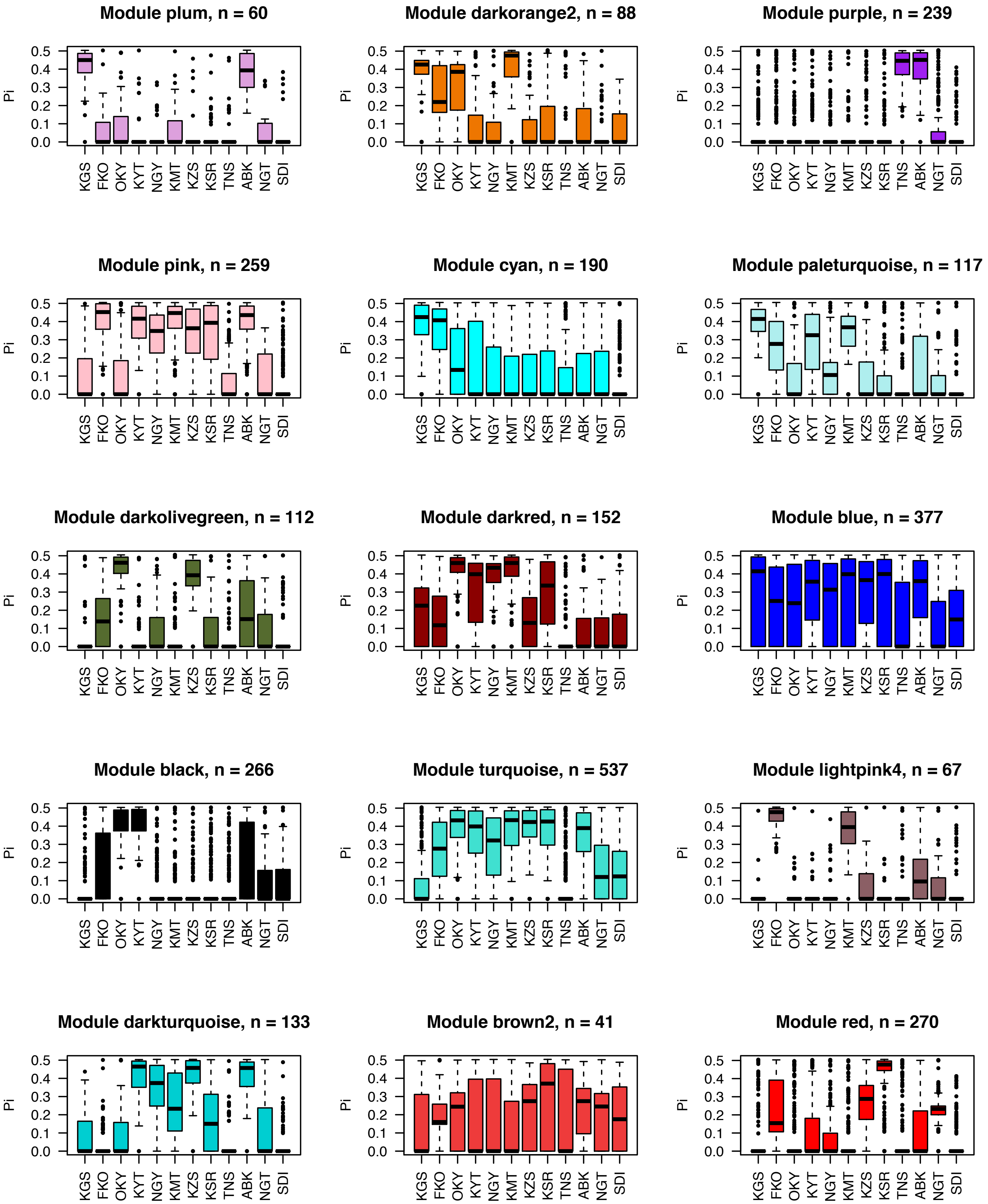


**Supplementary Figure S2** Nucleotide diversity of SNP modules across 12 natural populations of *Vicia sativa* from Japan.

Numbers of SNPs in each module are shown at the tops of boxplots. Three-letter codes indicate sampling location in Japan as shown in Supplementary Table S1 and Supplementary Figure S1.

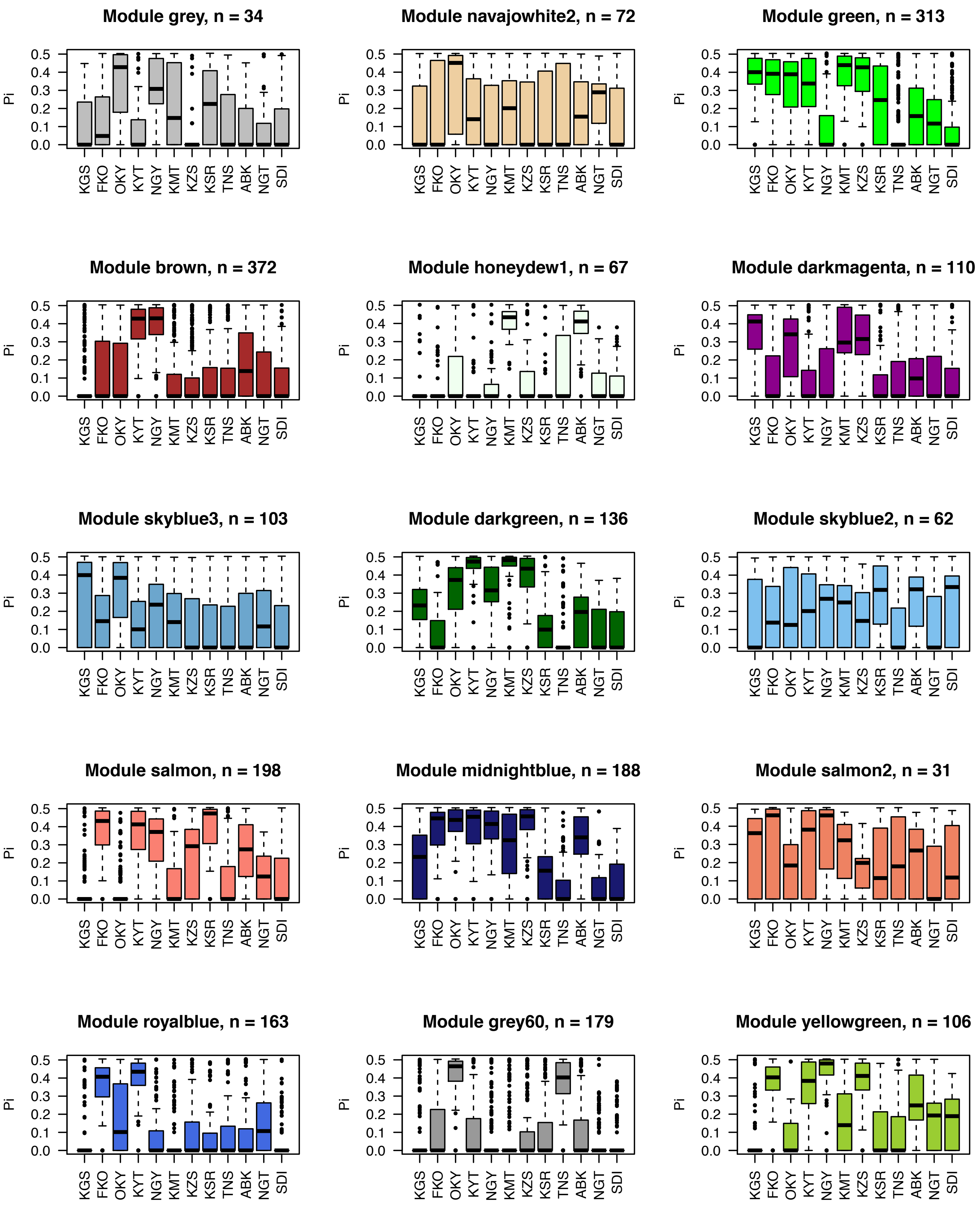


**Supplementary Figure S2** (continued)

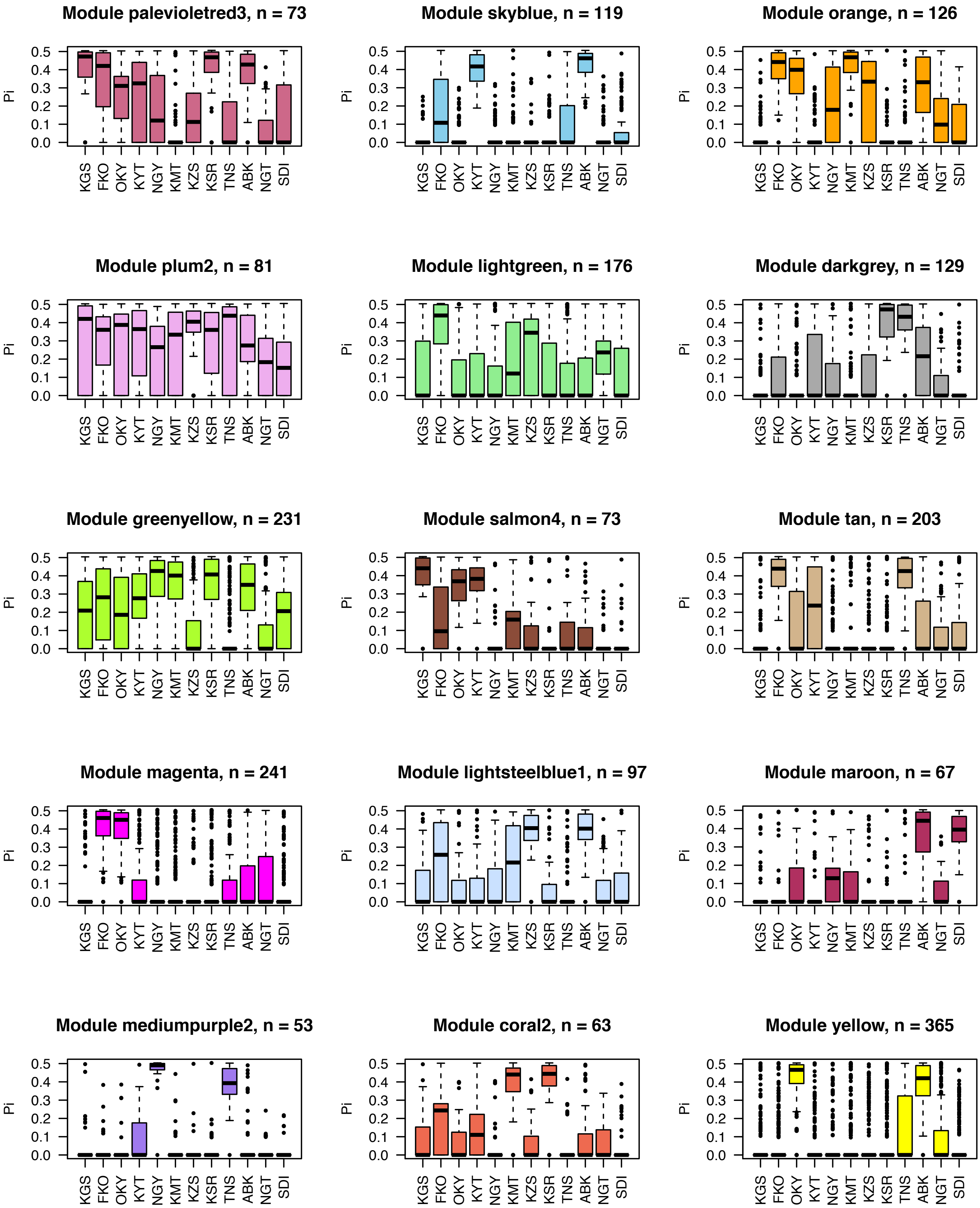


**Supplementary Figure S2** (continued)

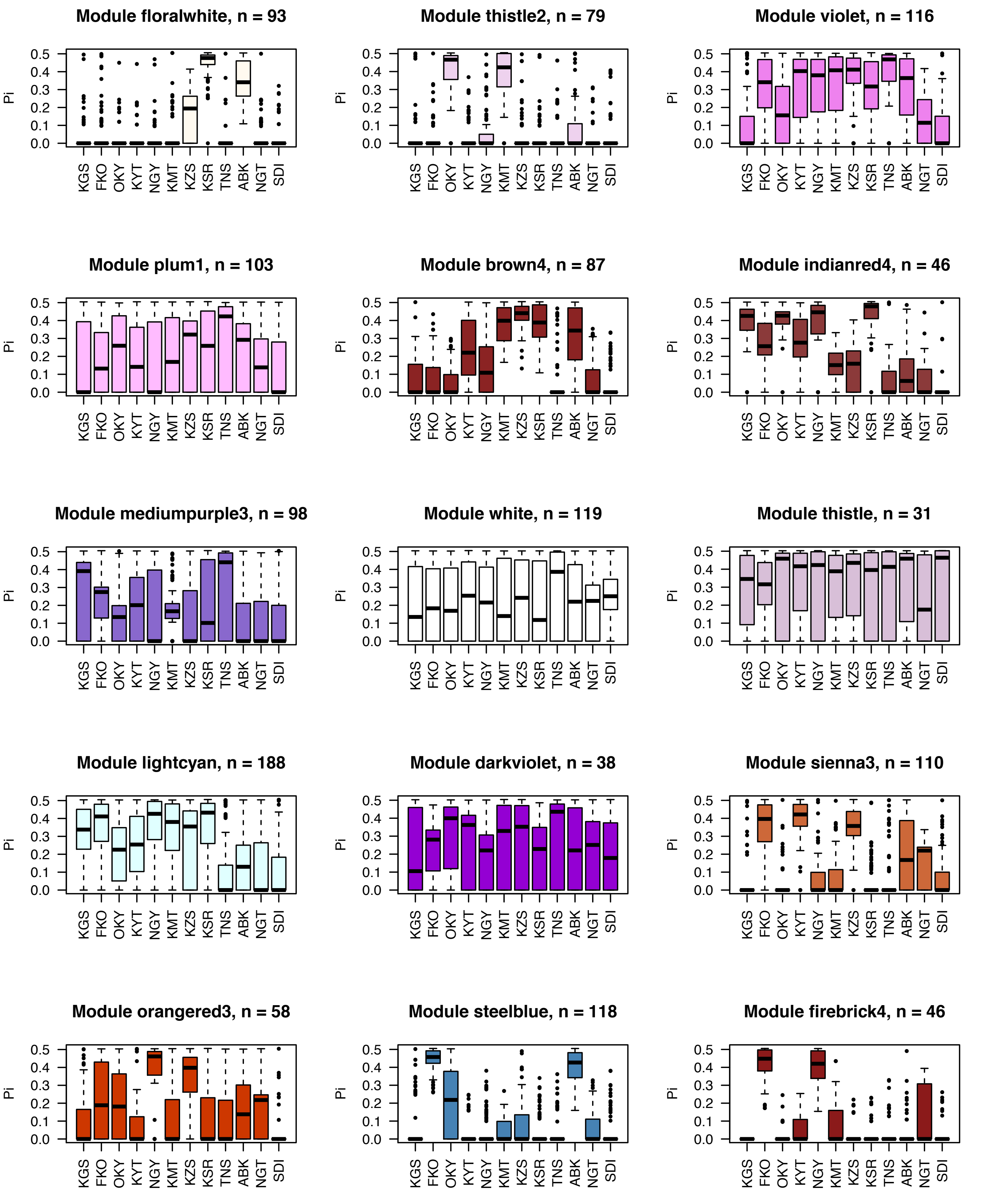


Supplementary Figure S2 (continued)

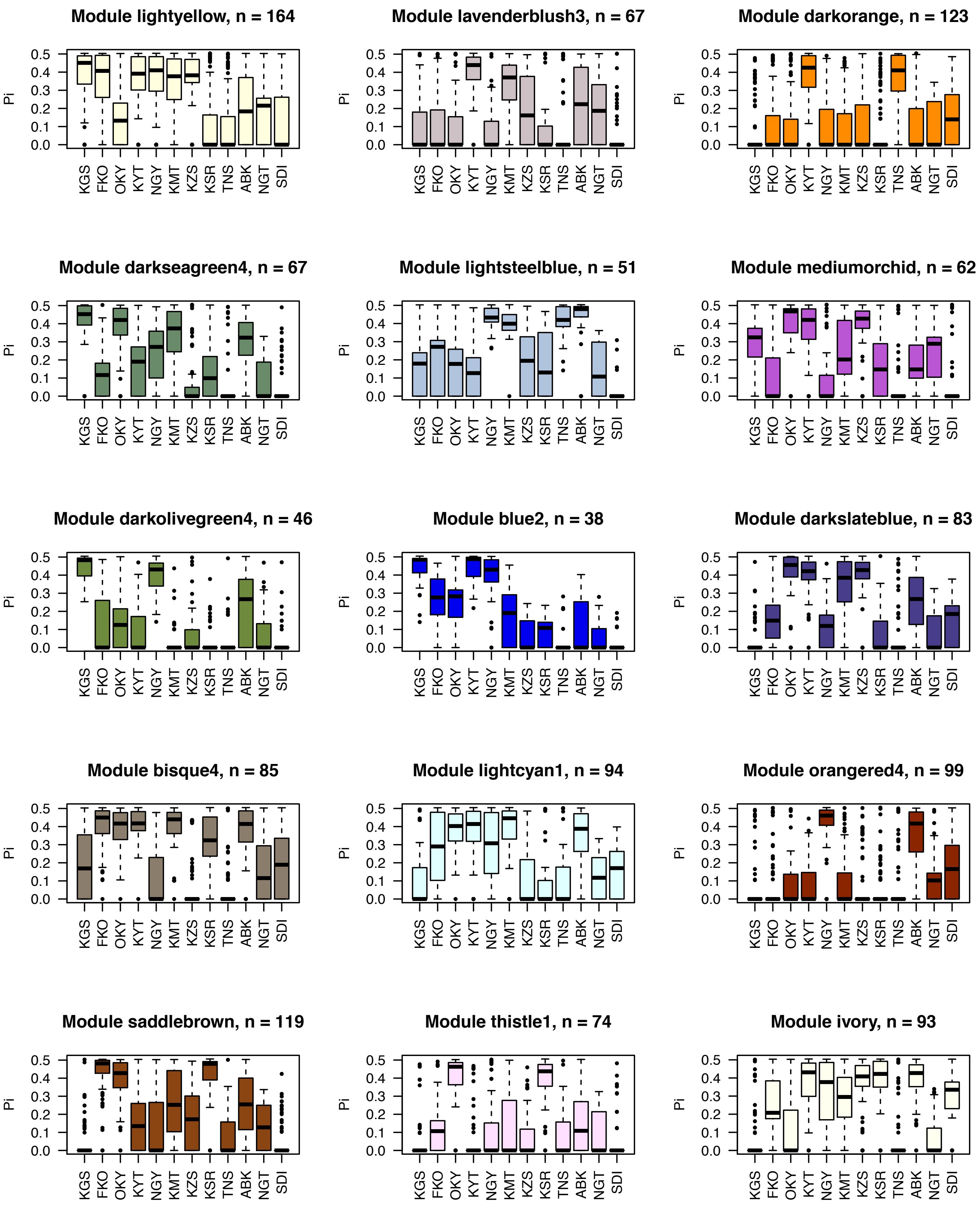


Supplementary Figure S2 (continued)

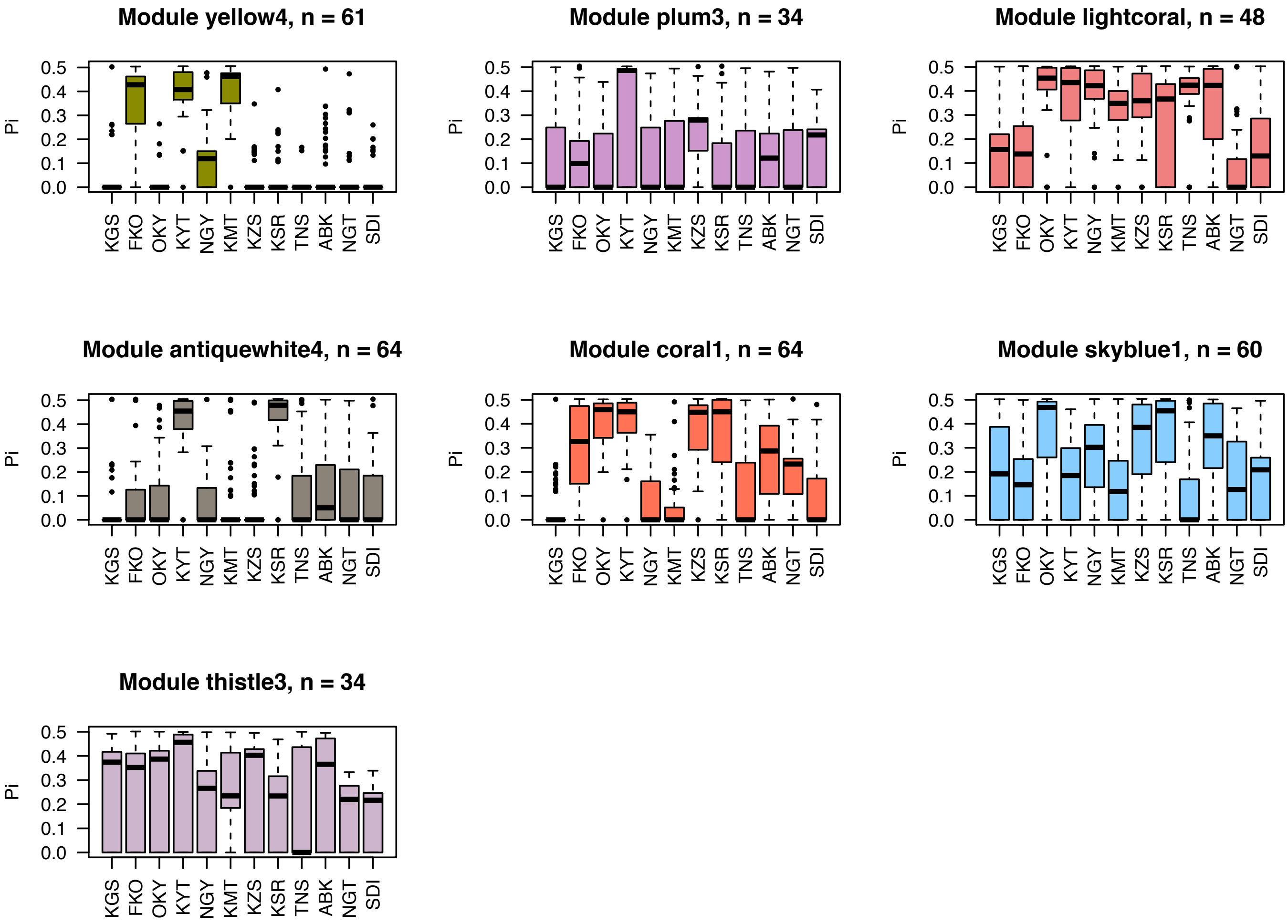


**Supplementary Figure S2** (continued)
